# Physiological Characterization under the Influence of Drought Stress and Salicylic Acid in *Valeriana wallichii* DC

**DOI:** 10.64898/2026.01.09.698547

**Authors:** Sadaf Ansari, Babita Patni, Devesh Jangpangi, Hem C. Joshi, Manoj Kumar Bhatt, Vijaykant Purohit

## Abstract

*Valeriana wallichii* DC, a high-altitude medicinal plant, is highly sensitive to drought stress, which adversely impacts its physiological functions and root biomass. This study aimed to evaluate salicylic acid’s role in mitigating drought stress by assessing physiological and biochemical responses in *V. wallichii*. Plants were treated with foliar applications of SA at concentrations of 0.25, 0.50, 0.75, and 1.0 mM during vegetative and flowering stages. Physiological parameters such as photosynthetic rate, stomatal conductance, water use efficiency (WUE), intrinsic WUE (iWUE), carboxylation efficiency, and biochemical traits including chlorophyll and carotenoid content were measured, alongside fresh and dry root biomass.Drought stress significantly reduced photosynthetic performance (Pn) (3.95 μmol CO_2_m□^2^s□^1^ at 25% FC), stomatal conductance, relative water content, and root yield. SA application notably improved these traits. At 25% FC, 1.0 mM SA enhanced WUE (28.78), iWUE (642.37 μmol/mol), and fresh root weight (8.41 g). Under moderate drought (50% FC), 0.50 mM SA improved net photosynthesis (12.5 μmol CO_2_ m□^2^s□^1^), dry root weight (2.31 g), and chlorophyll content. SA treatments also stabilized membrane integrity and maintained higher chlorophyll fluorescence efficiency (Fv/Fm).These findings demonstrate that foliar application of SA enhances drought tolerance in *V. wallichii* by improving photosynthetic efficiency, water relations, and root biomass. The study supports the strategic use of SA to improve the resilience and productivity of medicinal plants under drought stress conditions.

**Highlights of the study:**

1. Drought lowers growth and yield by reducing photosynthesis and water use.
2. Salicylic acid reduces drought effects and improves plant performance.
3. At severe drought, 1- 0.50 mM SA raised WUE, conductance, and root yield.
4. Under mild drought, 0.75- 0.50 mM SA improved gas exchange and yield.
5. SA doses 0.25-1 mM differ in effectiveness across drought stress stages.

## 1. Introduction

Climatic conditions and mobility among the seasons are one of the major defining factors for the sustainability of various ecosystems. An unexpected change in climate causes considerable interruption to both natural and agricultural ecosystems. It has been reported that the global temperature has increased rapidly in the last decade **(Sarma et al., 2024).** These adverse circumstances contribute to water deficit conditions that act as a major constraint to plant performance **(Irfan et al., 2023).** Drought is a meteorological phenomenon which is determined by a period without enough rain to cause a significant decline in soil moisture content by evapotranspiration and a decrease in plant growth as a consequence of physiological dehydration at a cellular level **(Li et al., 2023; Chen et al., 2022).** It affects plant physiology at all stages of development **(Takahashi et al., 2020; Miranda et al., 2023).** Physiological traits such as photosynthetic and gas exchange parameters are affected most by the decline in water levels **(Zhu et al., 2023).** Medicinal and aromatic plants are no exception to drought stress since they reduce herbal, essential oil and bio-active component yield. It reduces the absorption area of incident photosynthetic active radiation (PAR), decreases, efficiency of PSII and alters the stomatal closing and opening which plays role in the production and accumulation of primary and secondary metabolic compounds as well as dry matter content which ultimately reduces plant yield when the drought period coincides with critical growth period **(Moustakas et al., 2022; Nazari et al., 2023).** Plants response to stress is a complex process and involves numerous factors including signalling, transcription factors, hormones and secondary metabolites production. So, they have developed dynamic responses at morphological, physiological and biochemical levels allowing them to adapt different mechanisms for survival under unfavourable environmental conditions. Salicylic acid (SA) is an endogenously produced plant growth regulator which acts in plant defence response to abiotic stresses such as water and temperature stress **(Arif et al., 2020).** It is a vital signalling molecule that regulates plant growth and developmental processes via modulating signalling networks associated with biosynthesis pathways of plant secondary metabolites **(Song et al., 2023).** It renders plant resistance and tolerance to stresses and promotes plant growth. SA also influence physiological processes such as germination, stomatal closure, photosynthetic rate, transpiration, pigment content, membrane permeability, water uptake, nutrient uptake and stomatal conductivity **(Arikan et al., 2023).** Foliar spray of SA exerts control over stomatal opening and increases the photosynthetic rate and growth of the plant **(Altaf et al., 2023; Kusvuran & Yilmaz, 2023).** SA increases the chlorophyll content and affects the assimilation of CO_2_ and net absorbed radiation. This increase in the initial fluorescence (Fo), maximum primary productivity of PS II (Fv/Fo) and maximum quantum yield (Fv/Fm). An increase in these parameters under stress indicates the tolerance of plants which signifies the role of SA in drought tolerance **(Luqman et al., 2023; Feng et al., 2023) (Please insert Figure 1).**

**Fig. 1.**
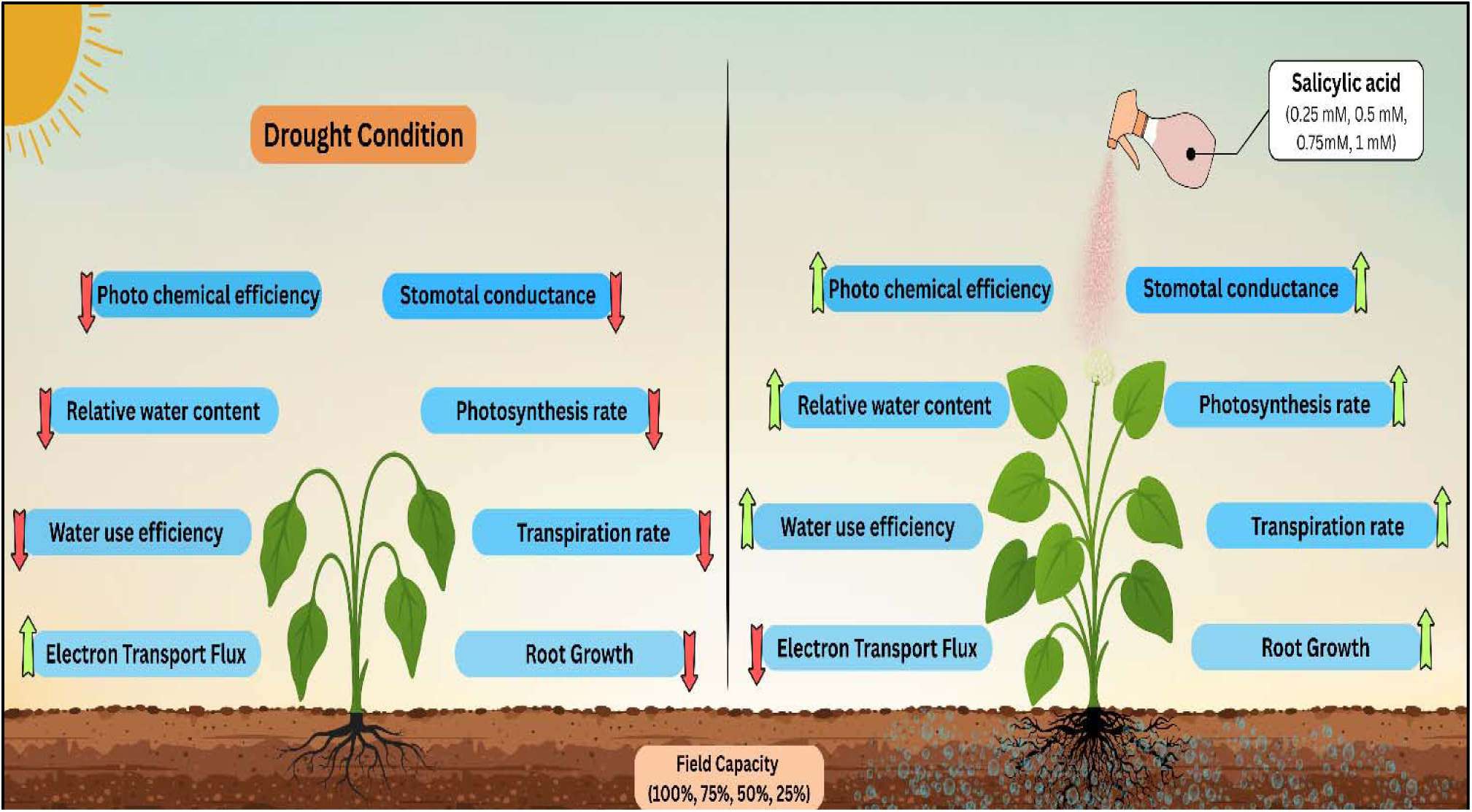
Pictorial representation of the effect of salicylic acid on *V. wallichii* under drought stress.

Medicinal and aromatic plants are important economic products which represent significant sources of economic revenue, foreign exchange and are among the most important agricultural export products. One of them, *Valeriana wallichii,* is a traditional medicinal plant **(Please insert Figure 2)** largely used in folk medicine. It has been reported to cure several ailments; thus, it is one of the key plants in the development of advanced herbal drugs. In *Valerian*, about 150 compounds of medicinal potential have been identified so far. Major bio-active constituents of plants belong to terpenes. Its product includes essential oil obtained from roots that are rich in iridoids, flavonoids, alkaloids, amino acids and lignanoids **(Jugran et al., 2019).** Roots also serve as the main source of therapeutic agents with spasmolytic, antibacterial and sedative properties.

**Fig. 2.**
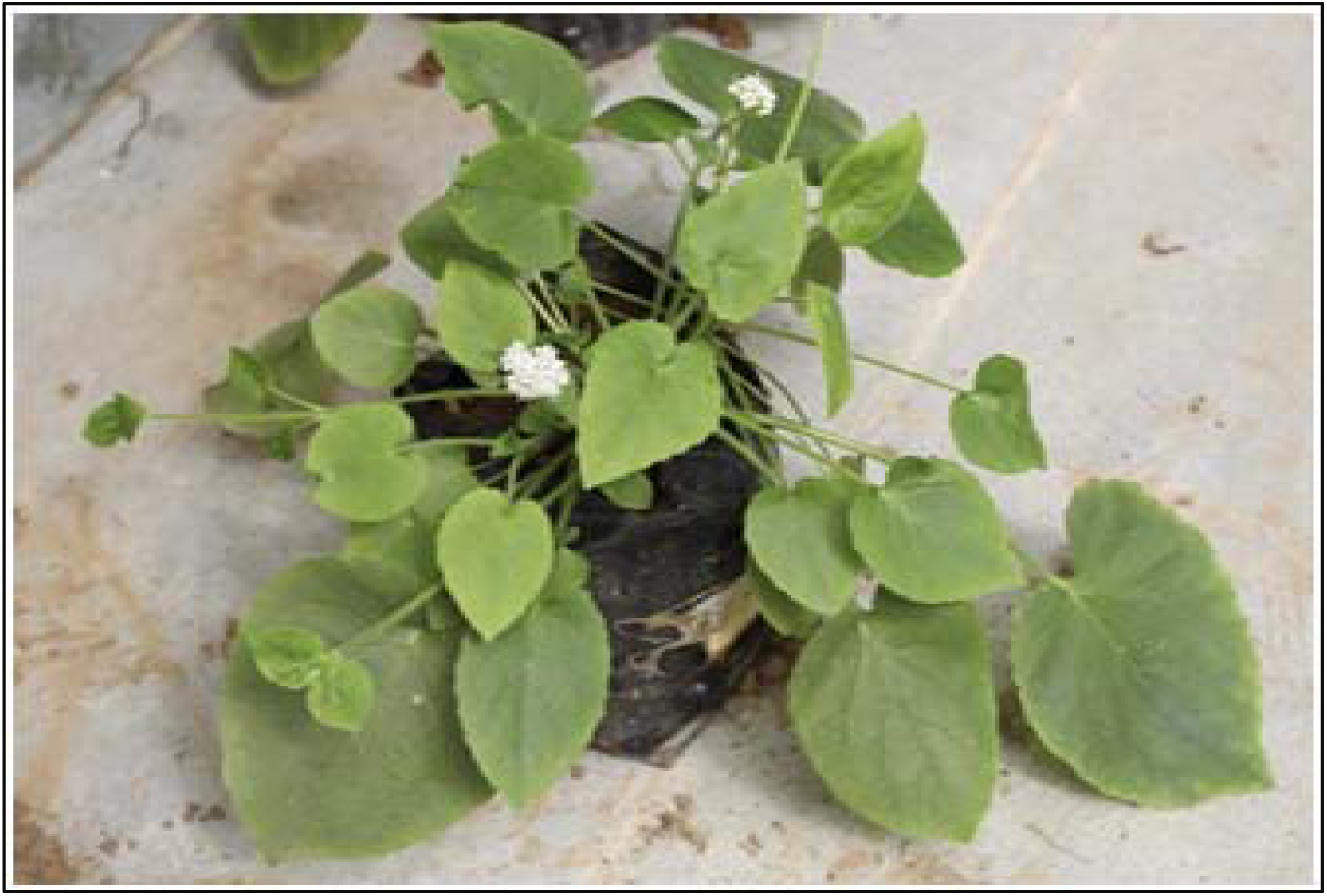
Valeriana wallichii.

Despite growing evidence of SA’s role in stress mitigation, the exact physiological mechanisms by which it enhances drought tolerance in medicinal plants remain poorly understood. Therefore, this study aims to investigate the drought-induced responses in *V. wallichii* and evaluate the impact of exogenous SA application on key physiological and root biomass traits. As an elicitor, SA offers a reliable, cost-effective, and accessible means to mitigate environmental stress. When applied at appropriate concentrations and developmental stages, SA may significantly enhance drought acclimation, supporting improved resilience and productivity in medicinal plants under changing climatic conditions.

## 1. Materials and Methods

### 2.1 Experimental site, climatic conditions and treatments detail

The present experiment was conducted under semi-controlled conditions at the Department of HAPPRC, H.N.B. Garhwal University (A Central University), Srinagar-Garhwal, Uttarakhand (India) 30°13’06” N 78°47’24”E at 700m above mean sea level during the years 2018-19 and 2019-20.Before applying drought stress, the plants were grown under semi-controlled glasshouse conditions using a standardized potting mixture of soil, sand, and compost. During the year 2018-19, the maximum temperature range between 11.56°C to 38.12°C and minimum temperature range between 9.09°C to 34.66°C and average maximum relative humidity was recorded 76.82% and average minimum relative humidity 66.32% and during the year 2019-20, the maximum temperature range between 8.58°C to 37.22°C and minimum temperature range between 7.54°C to 32.67°C and average maximum relative humidity was recorded 80.44% and average minimum relative humidity 72.01%. All plants were uniformly maintained at 100% field capacity for 1 month to allow for uniform establishment and acclimatization before drought and salicylic acid (SA) treatment, eliminating pre-treatment variability. One year oldrhizomes of *Valeriana wallichii* DC. (Planting material) were collected from the research field station Tala, Rudraprayag, Uttarakhand (India) (30°30’34”N 79°09’41”E), where plants were raised from seeds to ensure plant homogeneity. Uniformly sized and healthy rhizomes were carefully selected and washed to remove soil particles and foreign materials, followed by surface sterilization using Tween 20 and then transplanted in to the poly bags of size (20 × 20cm) and of 2.5litre capacity, which were filled with a prepared combination of soil, sand and dry plant litter (leaves) compost in a 2:1:1 ratio and maintained under semi-controlled glasshouse conditions. The water stress is given to plants at three levels after determining the field capacity and water-holding capacity of the polybag soil. There was a total of 20 treatments along with the control, each treatment was replicated thricely and each replicate contained 5 plants. The irrigation was scheduled in four different levels 100% field capacity (FC), 75% FC, 50% FC and 25% FC. The irrigation interval was determined based on the soil’s field capacity and water-holding capacity, ensuring optimal moisture availability for plant growth. It was scheduled weekly in winter and on alternate days in summer, aligning with seasonal evapotranspiration demands and simulating realistic drought conditions. Salicylic acid stock solution was prepared and using the standard stock solution 1mM, 0.75mM, 0.50mM and 0.25mM concentrations were prepared. Salicylic acid was applied as a foliar spray once daily for three consecutive days during the vegetative stage and the flowering initiation stage. The flowering stage application began at the visible onset of flowering.

### 2.2 Biochemical, Physiological estimations and root biomass

The observations were recorded after 15 days of foliar spray. For uniformity, fully expanded leaves from the main stem (simple leaves, not trifoliate) were collected during both the vegetative and flowering stages for all physiological and biochemical estimations.

### 2.3 Biochemical estimations Chlorophyll fluorescence parameters

Chlorophyll fluorescence was recorded using a plant efficiency analyzer (Handy PEA, Hansatech, UK) to assess PS II photochemical efficiency. Fully expanded third leaves were dark-adapted for 30 minutes using leaf clips before exposure to a saturating light pulse (3500 µmol photons m□^2^s□^1^). Fluorescence parameters such as Fo (initial fluorescence), Fm (maximum fluorescence), Fv (variable fluorescence), and Fv/Fm (maximum quantum yield of PS II) were calculated using the JIP test (Strasser and Strasser, 1995). Measurements were conducted between 09:00 to 11:00 AM on clear days at two growth stages, vegetative and reproductive to monitor PS II performance during development and stress conditions.

#### Chlorophyll Content

To measure the chlorophyll content following the method of **(Hiscox and Israelstam, 1979),** Fresh leaf samples (100 mg) were cut into fine pieces and placed in test tubes containing 10 ml of DMSO. Tubes were incubated at 60°C for 3–4 hours. After cooling to room temperature, the extracts were filtered and absorbance was measured at 645 nm and 663 nm using a UV-VIS spectrophotometer. The amount of chlorophyll was calculated by formula given by Arnon (1949).

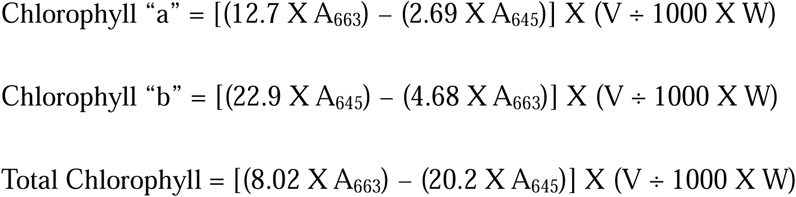

#### Carotenoid content

To measure the carotenoid content following the method of **(Hiscox and Israelstam, 1979),** 100mg of freshly cut fine pieces of leaf sample were taken in test tube containing 10ml of DMSO and incubated at 60°C for 3-4hrs. After incubation the tubes were taken out and cooled at room temperature. The extract was filtered and absorbance was recorded at 663nm, 645nm and 480nm. The amount of carotenoid was calculated by formula given by Arnon (1949).

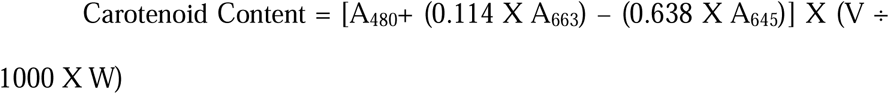

### 2.4 Root Biomass estimation Fresh root weight

Fresh weight was recorded with the help of digital weighing balance and expressed in grams. The fresh weight was measured immediately after harvest.

#### Dry root weight

Dry weight was recorded with the help of a digital weighing balance and expressed in grams. The dry weight was measured for three consecutive days until it remained constant after drying at 40°C.

### 2.5 Physiological estimation Relative water content (RWC)

To measure RWC following the method of **(Barrs and Weatherlay, 1962)**, the fully matured 4-5 leaves were collected at 12.00-13.00 hours and kept in separate plastic bags in the ice box. The fresh weight (FW) of collected leaf samples was recorded instantly after collection. The samples were dipped in de-ionized water in a closed aseptic Petri dish for a period of 12 hours until they attained full turgidity at room temperature and normal light conditions. After that, leaves were removed from the water and the turgid weight (TW) of the leaves was recorded after blotting them gently with blotting papers. The leaves were kept in paper packets and dried in a hot air oven at about 80°C until theyattained constant dry weight (DW).The Relative Water Content (RWC) was measured thricely (morning, afternoon and evening) after 15 and 30 days of treatments during vegetative and reproductive stages. The data were pooled for both vegetative and reproductive stages before analysis.

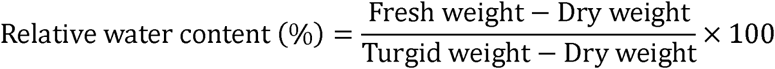

#### Membrane injury index

To measure the membrane injury index following the method of **(Chaisompogopan *et al.,* 1990),** the Leaf sample (0.1g) is cut into pieces of uniform size and taken in test tubes (two sets of 4 each) containing 10ml of double distilled water. One set of tubes is kept at 40-45°C for 30 minutes and another set at 100°C for 10-15 minutes in a water bath. After termination of treatment, samples are allowed to attain room temperature and the electrical conductivity of water in both sets of test tubes is recorded. The MSI was measured thricely (morning, afternoon and evening) after 15 and 30 days of treatments during the vegetative and reproductive stages. This was done to assess diurnal variation and the progression of membrane stability under different treatment conditions. The data were pooled for both vegetative and reproductive stages before analysis.

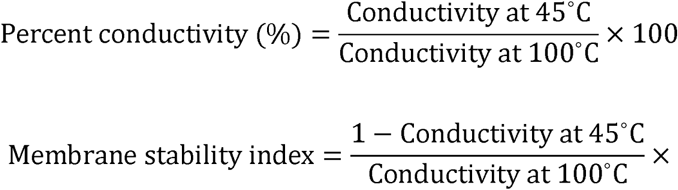

#### Leaf gas exchange parameters

Leaf gas exchange parameters were measured using a portable photosynthesis system (LI-6400XT, LICOR Inc., USA) under controlled conditions: photosynthetic photon flux density (PPFD) of 1600 µmol m□^2^s□^1^, CO□ concentration of 400 µmol mol□^1^, leaf temperature of 30°C, flow rate of 500 µmol s□^1^, and ambient relative humidity of ∼70%. Irradiance was provided by a red and blue LED light source. Flag leaves were placed in the leaf chamber until steady-state values were reached. Measurements were conducted between 9:00 to 11:00 AM on clear days at two growth stages vegetative and reproductive. At each growth stage, measurements were taken over a period of three consecutive days to ensure data reliability and account for validation of the results. Measurements included Photosynthetic rate (Pn) (µmol CO□m^−2^s^−1^), stomatal conductance (gs) (mol H□O m^−2^s^−1^), transpiration rate (Tr) (mmol H□O m^−2^s^−1^) and internal CO_2_ (Ci) (µmol CO□ mol^−1^). From these, additional parameters were calculated:

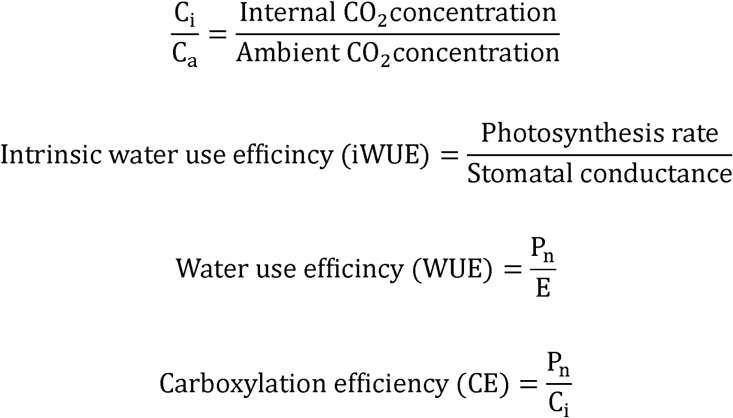

Where,

P_n_: Photosynthesis rate

C_i_: Internal CO_2_ concentration

E: transpiration rate

Drought tolerance is directly linked with the water status of the plants hence, to understand the drought tolerance mechanism, the determination of water related traits like RWC, MSI, electrolyte leakage, and electrical conductivity is studied during the experiment.

### 2.5 Statistical analysis

The data were analysed using R Studio (version 4.2.0), SPSS v25.0, and Microsoft Excel 2019. A completely randomized design (CRD) with three replications was used for all treatments. Two-way analysis of variance (ANOVA) was performed to determine significant differences among treatments at a 5% significance level (P ≤ 0.05). Mean comparisons were conducted using Duncan’s Multiple Range Test (DMRT) in R Studio. Standard error (SE) and critical difference (CD) were calculated to assess variability. Regression analysis and Pearson correlation were performed in R Studio to evaluate relationships between salicylic acid (SA) concentration and physiological responses. Principal Component Analysis (PCA) was used to identify key physiological traits influencing drought tolerance.

## 3. Results

### 3.1 Physiological Traits

#### Maximum photochemical efficiency (Fv/Fm)

Under drought stress conditions, Fv/Fm values declined notably at both vegetative and flowering stages **(Please insert Table 1)**. The most significant reduction was observed at 25% FC in the vegetative stage 0.595±0.037, while in the flowering stage, it decreased to 0.735±0.041 with 0.25mM SA. Foliar application of SA significantly improved Fv/Fm values under stress. Specifically, 0.75mM SA enhanced Fv/Fm to 0.789±0.008 at 25% FC during the vegetative stage and up to 0.825±0.001 at the flowering stage. Similarly, at 50% FC, 0.50mM and 0.25mM SA increased Fv/Fm to0.788±0.013and 0.793±0.014, respectively, indicating a dose-dependent ameliorative effect.

**Table 1.**
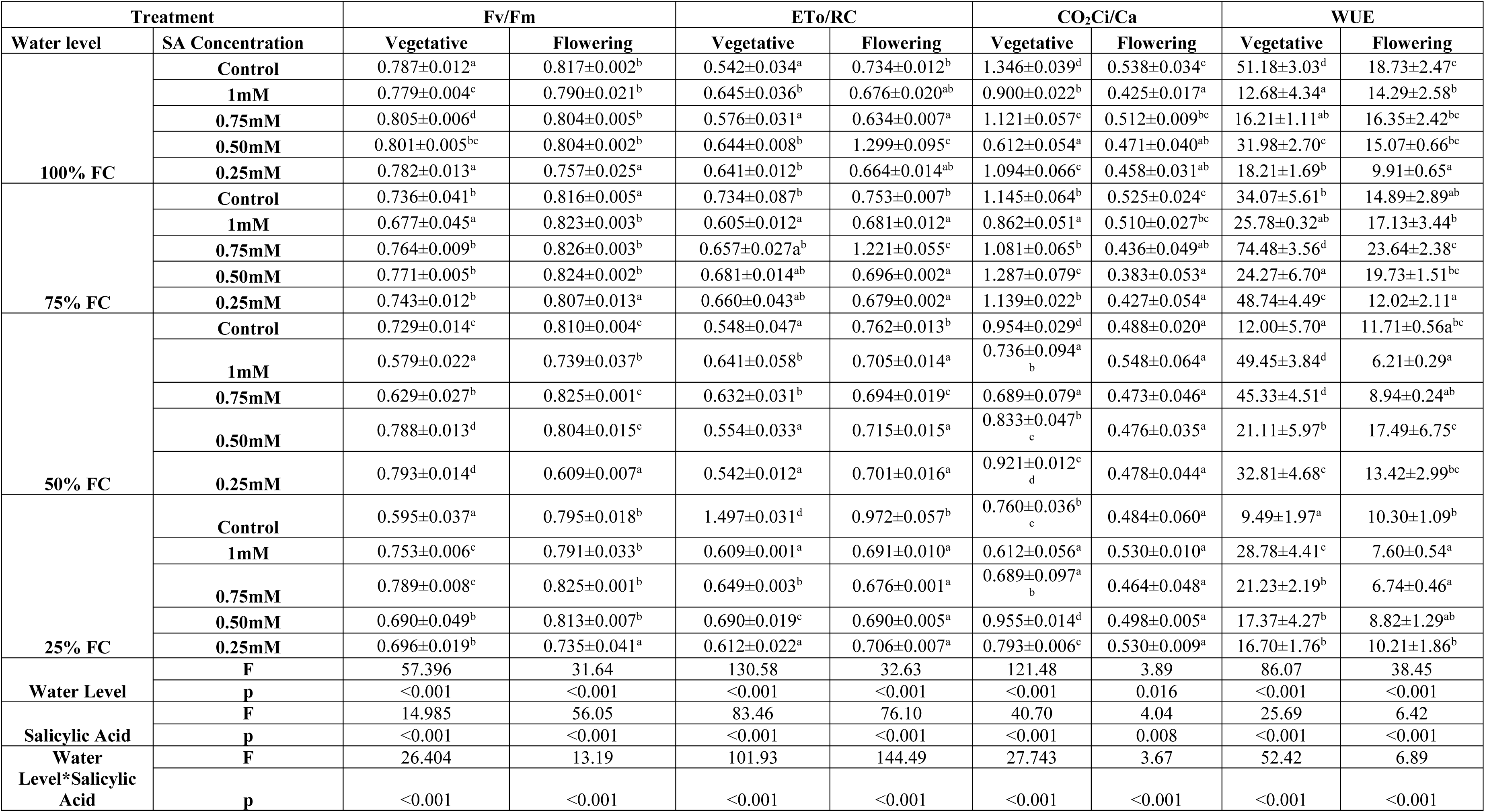
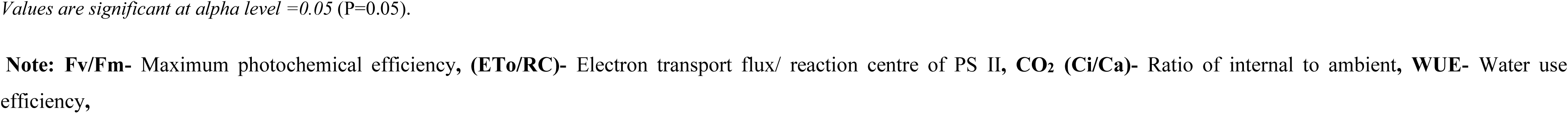
Effect of different concentration of foliar application of SA on physiological traits of *V. walliichii* at vegetative and flowering stage under various drought stress levels. ± indicates SE(m). Similar alphabets in the column showed non-significant deference.

#### Electron Transport flux per reaction centre of PS II (ETo/RC)

ETo/RC flux increased markedly with drought intensity, reaching the highest values at 25% FC in vegetative (1.497±0.031) and flowering stages (0.972±0.057) **(Table 1)**. Foliar SA application notably mitigated these effects. At 25% FC, 0.75mM SA reduced ETo to 0.649 in vegetative and 0.676±0.027 in the flowering stage, while 1mM SA also showed a significant reduction. At moderate stress (50% FC), 0.50mM SA was effective in reducing ETo to 0.554±0.033 during vegetative and 0.715±0.015 in flowering. Across all stress levels, higher SA concentrations (0.75mM, 1mM) consistently reduced ETo/RC values, reflecting better stress mitigation.

#### CO_2_ Internal to Ambient Ratio (Ci/Ca)

Drought stress significantly affected Ci/Ca ratios **(Table 1)**. The control plants at 25% FC had the lowest Ci/Ca (0.760±0.036 vegetative, 0.484±0.060 flowering). Foliar application of 0.50mM SA markedly increased Ci/Ca to 0.955±0.014 at 25% FC vegetative stage, showing a substantial improvement in CO_2_ assimilation efficiency. Similarly, under 75% FC, Ci/Ca was elevated to 1.287±0.079 with 0.50mM SA in vegetative stage. The ameliorating effect was dose-dependent, with moderate SA concentrations (0.50mM, 0.75mM) showing more pronounced improvements than the highest (1mM) concentration.

#### Water Use Efficiency (WUE)

WUE values were adversely affected under drought stress **(Table 1)**. However, foliar application of SA significantly improved WUE across stages. At 75% FC, 0.75mM SA increased WUE to 74.48±3.56 in vegetative and 23.64±2.38 in the flowering stage, showing the most substantial improvement. In the severe drought condition (25% FC), SA treatments elevated WUE from a control value of 9.49±1.97 to 21.23±2.19 with 0.75mM SA in the vegetative stage. Moderate drought stress (50% FC) showed enhanced WUE with 0.25mM and 0.50mM SA applications, recording values of 32.81±4.68 and 21.11±5.97, respectively.

#### Carboxylation Efficiency (CE)(μmolm^-2^s^-1^)

CE was significantly influenced by drought and SA treatments **(Please insert Table 2)**. The lowest CE was observed in control plants at 50% FC (1.186μmolm^-2^s^-1^vegetative) and 25% FC (4.30μmolm^-2^s^-1^flowering). Foliar SA application effectively improved CE. At 25% FC, 0.50mM SA enhanced CE to 5.76μmolm^-2^s^-1^ in the vegetative stage. Similarly, 0.75mM SA showed improvement at 75% FC, increasing CE to 5.93μmolm^-2^s^-1^ vegetative. In flowering stages, CE was highest at 100% FC with 1mM SA (6.34μmolm^-2^s^-1^). These findings suggest that moderate SA concentrations ameliorate the negative effects of drought on carboxylation capacity, likely through improved Rubisco activity and CO□ assimilation. CE was significantly influenced by drought and SA treatments **(Table 2)**. The lowest CE was observed in control plants at 50% FC (1.18μmolm^-2^s^-1^vegetative) and 25% FC (4.30μmolm^-2^s^-1^ flowering). Foliar SA application effectively improved CE. At 25% FC, 0.50mM SA enhanced CE to 5.76μmolm^-2^s^-1^ in the vegetative stage. Similarly, 0.75mM SA showed improvement at 75% FC, increasing CE to 5.93μmolm^-2^s^-1^ vegetative. In flowering stages, CE was highest at 100% FC with 1mM SA (6.34μmolm^-2^s^-1^). These findings suggest that moderate SA concentrations ameliorate the negative effects of drought on carboxylation capacity, likely through improved Rubisco activity and CO□ assimilation.

**Table 2:**
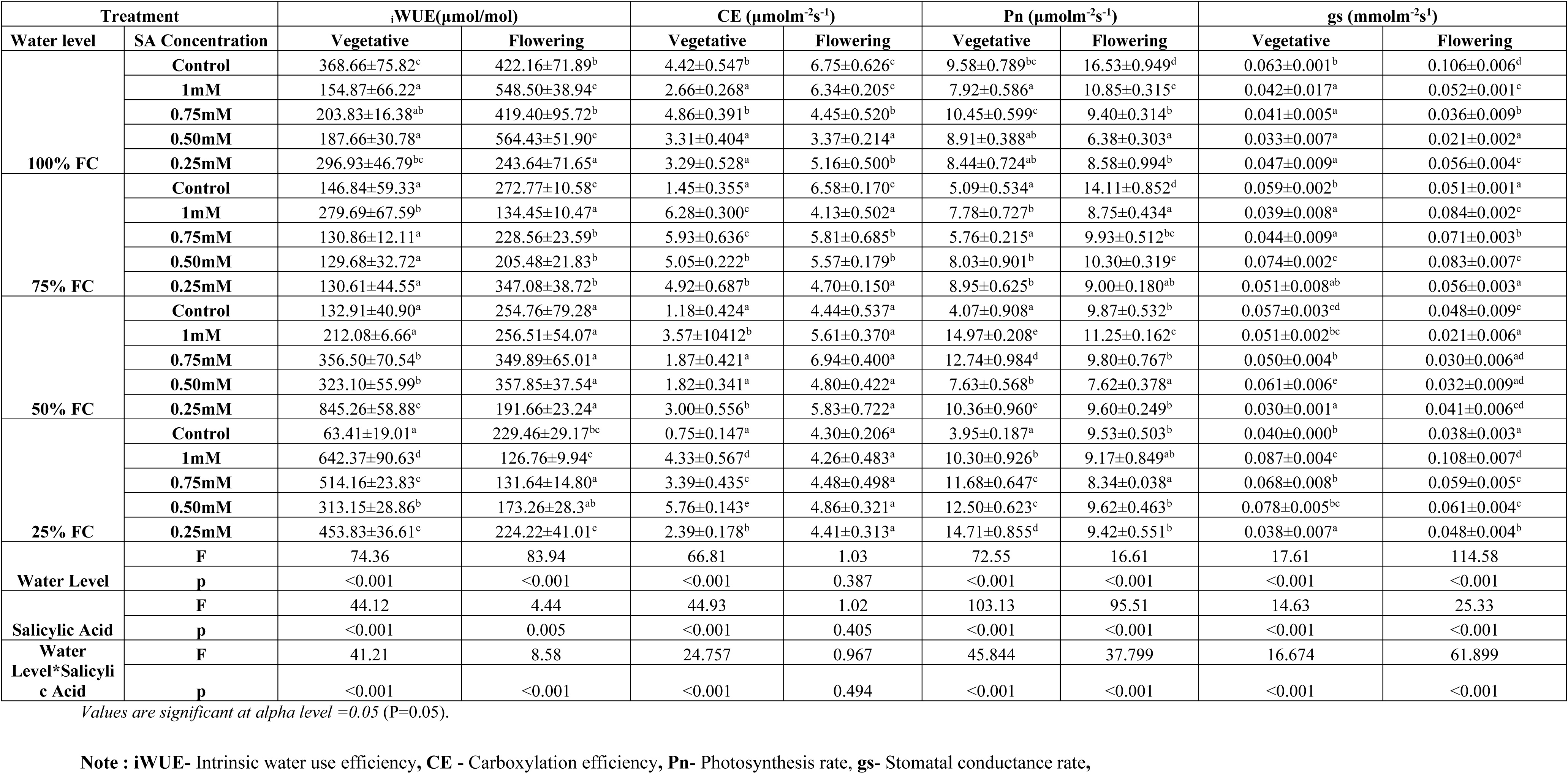
Effect of different concentration of foliar application of SA on physiological traits of *V. walliichii* at vegetativeand flowering stage under various drought stress levels. ± indicates SE(m).Similar alphabets in the column showed non-significant deference.

#### Net Photosynthetic Rate (Pn) (μmolm^-2^s^-1^)

A significant reduction in Pn was recorded under drought stress, especially in control plants at 25% FC (3.95μmolm^-2^s^-1^ vegetative, 9.53μmolm^-2^s^-1^ flowering) **(Table 2)**. SA foliar treatments notably enhanced Pn values. In the vegetative stage, 1mM SA increased Pn to 14.97μmolm^-2^s^-1^ at 50% FC, while at 25% FC, 0.50mM SA improved Pn to 12.50μmolm^-2^s^-1^. Flowering stage showed significant improvements at 1mM SA (11.25μmolm^-2^s^-1^ at 50% FC) and 0.50mM SA (10.30μmolm^-2^s^-1^ at 75% FC). These observations highlight the potential of SA in maintaining photosynthetic rates under limited water availability.

#### Stomatal Conductance (gs) (mmol^-2^s^-1^)

Stomatal conductance (gs) exhibited significant variations with water stress and SA application **(Table 2)**. The lowest gs was recorded at 100% FC with 0.50mM SA (0.033mmol^-2^s^-1^vegetative) and at 50% FC flowering (0.021mmol^-2^s^-1^) with 1mM SA. Under severe drought (25% FC), SA 1mM significantly increased gs to 0.087 in vegetative and 0.108 in flowering stage, suggesting a possible SA-induced stomatal opening. At moderate stress levels (75% FC), gs was enhanced by 0.50mM SA (0.074mmol^-2^s^-1^ vegetative, 0.083mmol^-2^s^-1^ flowering). These results indicate that SA modulates stomatal responses, contributing to better gas exchange under drought stress.

#### Intrinsic Water Use Efficiency (iWUE)

During drought stress, iWUE significantly fluctuated at both vegetative and flowering stages **(Table 2)**. Severe drought (25% FC) reduced iWUE to 63.41±19.01 in vegetative and 229.46±29.17in flowering stage in control plants. Foliar application of SA significantly enhanced iWUE. At 25% FC, 1mM SA treatment drastically increased iWUE to 642.37±90.63 in vegetative stage, while 0.75mM and 0.25mM SA also showed remarkable improvement (514.16±23.83 and 453.83±36.61, respectively). A similar trend was observed at 50% FC, where 0.25mM SA elevated iWUE to 845.26±58.88 during the vegetative stage. In the flowering stage, 0.50mM SA significantly improved iWUE at 100% FC (564.43±51.90). These results indicate that SA foliar sprays significantly improve water use efficiency under drought, with pronounced effects at moderate to severe stress levels.

#### Intercellular CO_2_ Concentration (Ci) (μmol mol^-1)^

Drought stress significantly affected Ci values at both vegetative and flowering stages **(Please insert Table 3)**. The lowest Ci was observed at 25% FC in vegetative stage with 1mM SA (247.42μmol mol^-1^) and at flowering stage with 0.50mM SA (162.33μmol mol^-1^). Foliar application of SA at 0.50mM notably reduced Ci at 100% FC (249.05μmol mol^-1^ vegetative, 199.56μmol mol^-1^ flowering). However, at 75% FC, 0.75mM SA decreased Ci values (394.73μmol mol^-1^ vegetative, 183.29μmol mol^-1^ flowering). These results indicate that SA application regulates stomatal behaviour and internal CO_2_ diffusion under drought, maintaining photosynthetic activity.

**Table 3:**
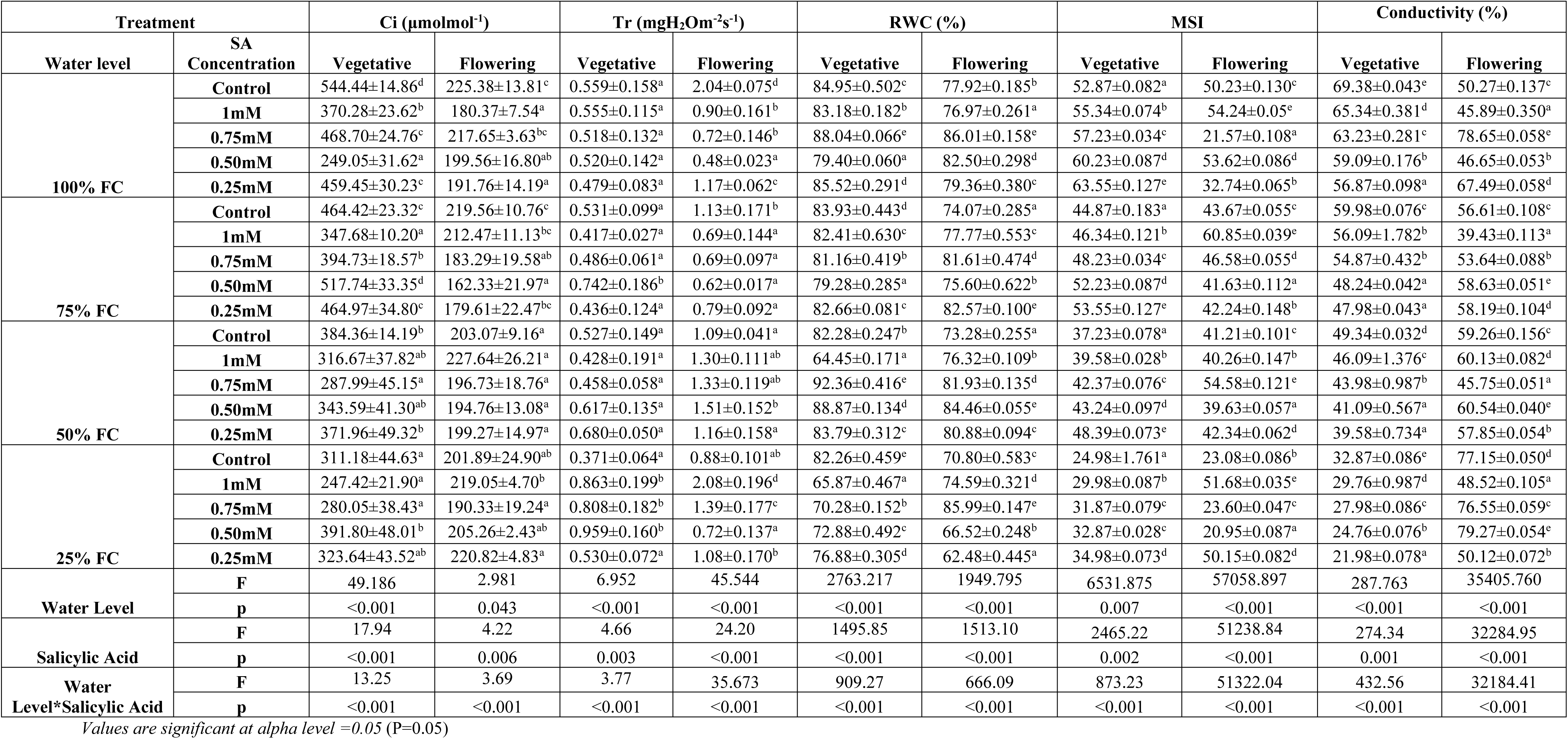
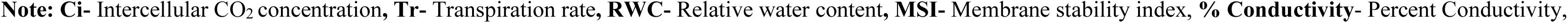
Effect of different concentration of foliar application of SA on physiological traits of *V. walliichii* at vegetativeand flowering stage under various drought stress levels. ± indicates SE(m).Similar alphabets in the column showed non-significant deference.

#### Transpiration Rate (Tr)(mgH_2_Om^-2^s^-1^)

Drought stress reduced transpiration rate (Tr), with the lowest values recorded at 100% FC with 0.25mM SA (0.479mgH_2_Om^-2^s^-1^ vegetative) and at 75% FC with 0.50mM SA (0.62mgH_2_Om^-2^s^-1^ flowering) **(Table 3)**. Under severe drought (25% FC), SA 0.50mM and 0.75mM significantly increased Tr (0.959mgH_2_Om^-2^s^-1^ and 0.808mgH_2_Om^-2^s^-1^ respectively in vegetative stage). Interestingly, at 25% FC, 1mM SA increased Tr to 0.863mgH_2_Om^-2^s^-1^ in vegetative stage and 2.08±0.196 in flowering stage. This suggests that SA application modulates plant water loss, potentially improving cooling and nutrient transport under stress.

#### Relative Water Content (RWC)

RWC significantly decreased under drought stress, with the lowest recorded at 50% FC in vegetative stage (64.45% with 1mM SA) and 25% FC in flowering stage (62.48% with 0.25mM SA) **(Table 3)**. However, 0.75mM SA application at 50% FC improved RWC to 92.36% in vegetative stage, while 0.75mM SA at 25% FC enhanced flowering stage RWC to 85.99%. These results confirm that SA plays a crucial role in maintaining cellular hydration and osmotic balance under drought stress conditions.

#### Membrane Stability Index (MSI)

MSI was notably decreased under severe drought (25% FC), with the lowest values recorded in vegetative stage control plants (24.98%) and flowering stage (20.95% with 0.50mM SA) **(Table 3)**. SA application improved MSI significantly. At 25% FC, 0.25mM SA increased MSI to 34.98% in vegetative stage and 50.15% in flowering stage. Similarly, at 100% FC, 0.25mM SA enhanced MSI to 63.55% vegetative. This indicates that SA application enhances membrane integrity, reducing drought-induced lipid peroxidation.

#### Percentage Conductivity (%)

Conductivity, a marker of membrane damage, increased under drought stress. The highest conductivity was observed at 100% FC with 0.75mM SA (78.65% flowering stage) and 25% FC with 0.50mM SA (79.27% vegetative stage) **(Table 3)**. However, lower conductivity was maintained with 0.25mM SA at 25% FC (21.98% vegetative stage). These results reflect the protective effect of SA in reducing membrane leakage under drought by enhancing antioxidant defenses.

### 3.2 Biochemical Traits Chlorophyll Content (mg/gFW)

#### Chlorophyll “a”

Chlorophyll “a”content was significantly affected by drought stress, with maximum reductions observed at 25% FC in both vegetative (6.375 mg/gFW) and flowering (6.621 mg/gFW) stages in control plants **(Please insert Table 4)**. Foliar application of SA ameliorated this decline. At 25% FC, 0.75mM SA increased chlorophyll “a” to 5.840 mg/g FW during vegetative and 1mM SA enhanced it to 6.608 mg/g FW in flowering. Similarly, at 50% FC, 0.50mM SA increased chlorophyll “a” to 6.783 mg/gFW (vegetative) and 6.924 mg/gFW (flowering). At 75% FC, 0.25mM SA showed the best effect, enhancing chlorophyll “a” to 5.832 mg/gFW (vegetative) and 7.135 mg/gFW (flowering). In well-watered conditions (100% FC), 0.50mM SA and 1mM SA were most effective during vegetative (6.685 mg/gFW) and flowering (6.841 mg/gFW) stages, respectively.

**Table 4:**
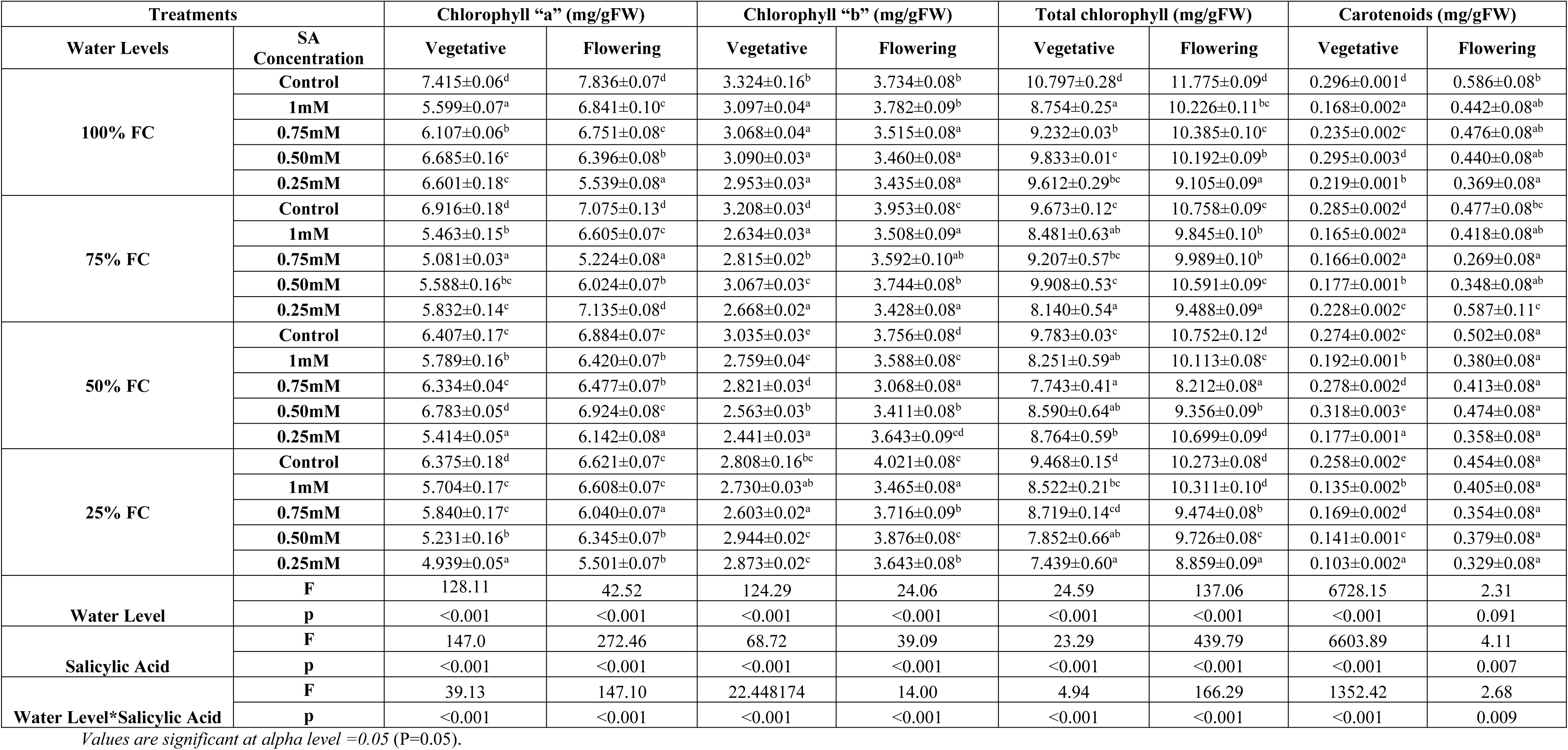
Effect of different concentration of foliar application of SA on biochemical traits of *V. walliichii* at vegetative and Vegetative stage under various drought stress levels. ± indicates SE(m). Similar alphabets in the column showed non-significant deference.

#### Chlorophyll “b”

Chlorophyll “b” content exhibited a decreasing trend with increasing drought stress. The lowest values were recorded at 25% FC in control plants. Foliar application of 0.50mM SA significantly improved chlorophyll “b” at 25% FC (2.944 mg/gFW vegetative, 3.876 mg/gFW flowering)**(Table 4).** At 50% FC, 0.75mM SA (2.821 mg/g FW vegetative) and 0.25mM SA (3.643 mg/gFW flowering) were effective. For 75% FC, 0.50mM SA showed maximum improvement (3.067 mg/gFW vegetative, 3.744 mg/gFW flowering). At 100% FC, 1mM SA enhanced chlorophyll “b” (3.097mg/gFW vegetative, 3.782mg/gFW flowering). These increments were significant compared to respective controls.

#### Total Chlorophyll

Total chlorophyll content declined under drought stress, reaching minimum values at 25% FC (Table 4). SA application significantly increased total chlorophyll content. At 25% FC, 0.75mM SA elevated it to 8.719 mg/gFW (vegetative), while 1mM SA raised it to 10.311 mg/g FW (flowering). At 50% FC, 0.25mM SA (8.764 mg/g FW vegetative) and 10.699 mg/g FW (flowering) showed the best results. Similarly, 0.25mM and 0.50mM SA at 75% FC increased total chlorophyll content to 9.908mg/gFW and 10.591 mg/gFW, respectively. Under 100% FC, 0.50mM SA (9.833 mg/gFW vegetative) and 0.75mM SA (10.385 mg/gFW flowering) were most effective. All SA treatments significantly enhanced total chlorophyll content compared to controls.

#### Carotenoids

Carotenoid content was significantly reduced under drought, especially at 25% FC in control plants (0.258 mg/gFW vegetative, 0.454 mg/gFW flowering) **(Table 4).** SA foliar application markedly improved carotenoid levels. At 25% FC, 0.75mM SA (0.169 mg/gFW vegetative) and 1mM SA (0.405 mg/gFW flowering) were effective. At 50% FC, carotenoid content increased by 0.50mM SA (0.318mg/gFW vegetative, 0.474mg/gFW flowering). At 75% FC, 0.25mM SA raised carotenoids to 0.228 mg/gFW (vegetative) and 0.587 mg/g FW (flowering). For 100% FC, 0.50mM SA enhanced carotenoid content to 0.295 mg/gFW in vegetative, while 0.75mM SA increased it to 0.476 mg/g FW during flowering. All increases were significant.

### 3.3 Root Biomass Traits

#### Fresh Root Weight (g)

Fresh root weight was significantly reduced under drought stress, with the lowest values observed at 25% FC (2.53 g vegetative, 1.09 g flowering) in control plants **(Please insert Table 5).** Foliar application of SA effectively mitigated these reductions. At 25% FC, 1mM SA significantly increased fresh root weight to 8.41 g (vegetative) and 9.56 g (flowering). Similarly, at 50% FC, 0.50mM SA enhanced fresh root weight to 9.46 g (vegetative) and 9.58 g (flowering). At 75% FC, 1mM SA also improved fresh root weight significantly (9.25 g vegetative, 7.48 g flowering). In well-watered plants (100% FC), 0.25mM SA was most effective, increasing fresh root weight to 3.62 g (vegetative) and 11.11 g (flowering). These improvements indicate SA’s significant role in alleviating drought-induced biomass reduction.

**Table 5:**
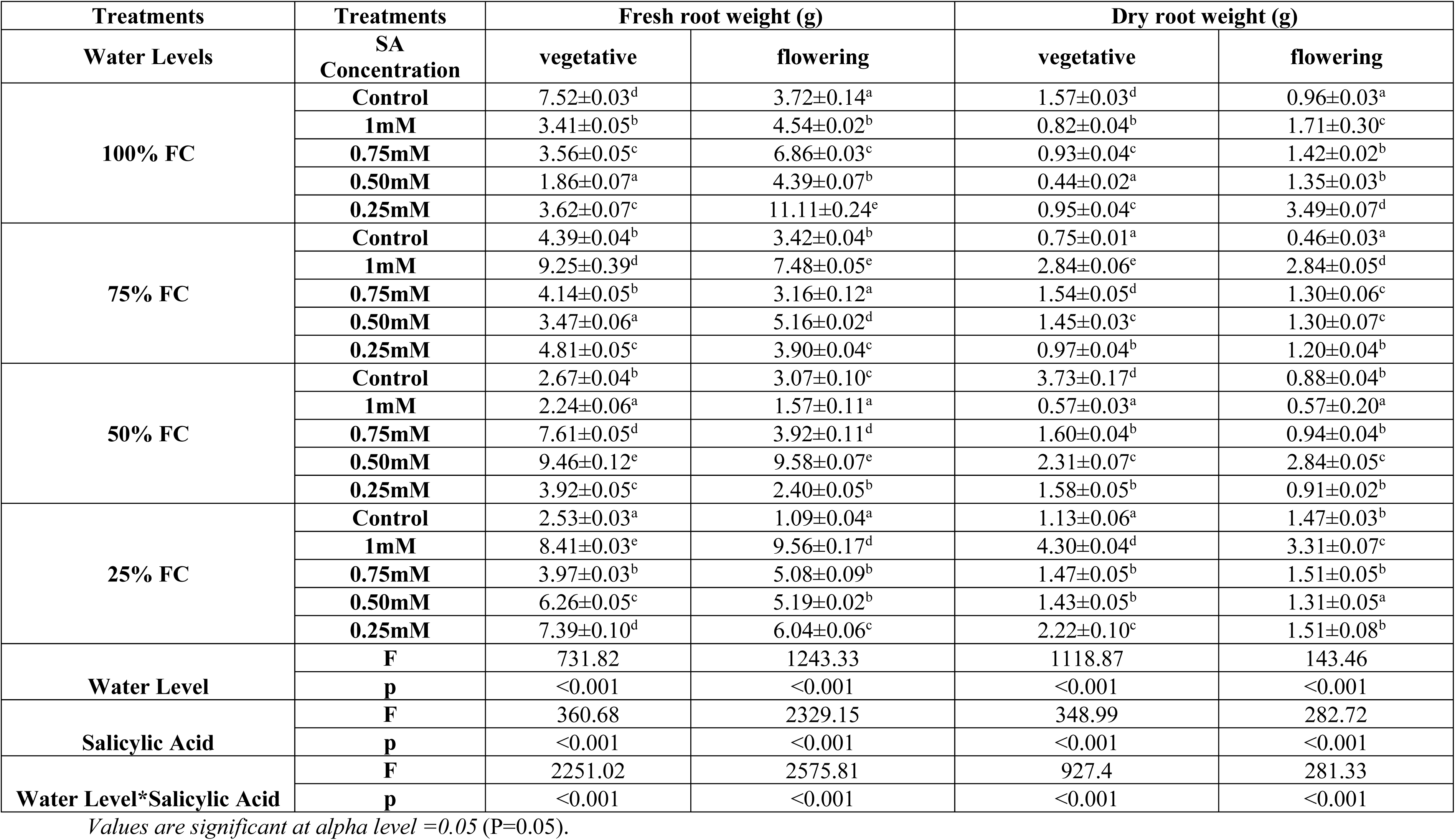
Effect of different concentration of foliar application of SA on root biomass traits of *V. walliichii* at vegetative and flowering stage under various drought stress levels. ± indicates SE(m). Similar alphabets in the column showed non-significant deference.

#### Dry Root Weight (g)

Dry root weight followed a similar trend, with maximum reduction in control plants at 25% FC (1.13 g vegetative, 1.47 g flowering) **(Table 5).** Foliar application of 1mM SA significantly enhanced dry root biomass at 25% FC to 4.30 g (vegetative) and 3.31 g (flowering). At 50% FC, 0.50mM SA treatment was most effective, increasing dry root weight to 2.31 g (vegetative) and 2.84 g (flowering). At 75% FC, 1mM SA again resulted in the highest dry root weight (2.84 g for both vegetative and flowering). Under 100% FC, 0.25mM SA improved dry root weight to 0.95 g (vegetative) and 3.49 g (flowering). These results suggest that SA enhances root biomass accumulation under drought by mitigating stress impacts on root growth.

## 4. Discussions

### 4.1 Physiological and Biochemical Traits

In recent times, drought has been considered an important factor in limiting crop growth and yield. Damage in photosynthetic apparatus inhibited maximum photosynthetic rate, light-use efficiency and increased respiration rate due to water shortage. Alterations in photosynthetic parameters under different levels of drought stress are mainly caused by stomatal and non-stomatal factors, which manifest by changes in C_i_, while C_a_ is relatively stable. Due to the closure of stomata, CO_2_ availability becomes low which shows alteration in carbon metabolism **(Hasanagi**ć **et al., 2020)**. The reduction in root water uptake under drought conditions shows rapid stomatal closure subsequently affects photosynthetic apparatus through declines in CO_2_ assimilation rate, transpiration and intercellular CO_2_ **(Qiao et al., 2024).** Gas exchange parameters, light assimilation rate and membrane stability were greatly affected by the drought in *Valeriana*. Photosynthetic traits are one of the major parameters that represent the plant growth. In the experiment with increasing drought stress levels: photosynthetic rate (P_n_), stomatal conductance (g_s_), transpiration rate (T_r_) and intercellular CO_2_ (C_i_) decrease significantly and similarly decline in water use efficiency (WUE), intrinsic water use efficiency (_i_WUE), carboxylation efficiency (CE) as well as ratio of C_i_ to C_a_ was observed as water level reduces to 25% FC from 100% FC. A similar response towards gas exchange parameters was observed in *Solanum lycopersicum* under drought stress **(Liang et al., 2020).** In *Solanum lycopersicum*, drought stress restricts plant growth by reducing photosynthesis, gaseous exchange and light assimilation rate. Accordingly, the successive decrease of g_s_ in the presence of high drought stress levels confirmed stomatal limitation for photosynthesis restriction activity compared to the well-watered plants **(Hasanagi**ć **et al., 2020).** The declining transpiration rate reflects an earlier plant response to a reduction in the water potential around the root zone. Plants showed a high ability to control the water fluxes through higher stomatal closure, which might be considered the first line of defence against water loss and protection mechanism against tissue dehydration **(Hayat et al., 2020).** The severity of water deficits led to a drastic reduction in water use efficiency. Adaptation in WUE by plants is synchronized by the association between carbon assimilation and water consumption, which is an imperative mechanism used by plants to withstand water shortage. Intrinsic water use efficiency is the ratio of CO_2_ assimilation to transpiration. It is the negative function of C_i_/C_a_. Stomatal control and morphological root architecture are considered the key controlling factors for WUE under drought stress **(Yang et al., 2021).** WUE expresses the relation between root water uptake ability and shoot water use. It is commonly recognized that species having higher WUE are better adapted to drought. The increase in water use efficiency under water deficit describes the fact that either the biomass production is reduced **((Blankenagel et al., 2018; Yu et al. (2019)** or the net CO_2_ assimilation rate is being less than transpiration rate **(Khatri & Rathore, 2019).** Alternatively, it was also possible that a partial inhibition of the photosynthetic function of CO_2_ fixation, RUBP carboxylation and inorganic phosphorous transformation occurred **(Flexas et al., 2016)**. These stomatal structural-functioning changes might help the plants to adjust effectively to unfavourable environmental conditions. The decrease in photosynthesis is due to the restricted gaseous exchange by closure of stomata, reduction in leaf area, water and nutrient absorption and consequently reduced cooling of a leaf by reduced transpiration rate causing damage to photosynthetic apparatus and cellular membrane **(Kapoor et al., 2020)**. **(Wang et al., 2016)** also reported a reduction in gas exchange parameters and yield due to increase in drought stress levels while working on wheat. The diminution in leaf gas exchange parameters might be due to the degradation of chlorophyll content and reduced nitrogen uptake.

Over the time, different studies suggested the role of SA in mitigating drought stress. Our findings confirm that foliar application of SA at different concentration has significantly increases the gas exchange parameters (*i.e.,* leaf stomatal conductance and CO_2_ assimilation rate, intercellular CO_2_, photosynthesis rate and transpiration rate) in plants under study. In *Valeriana* during vegetative stage, it was observed that at 25% FC, 0.25mM, 0.50mM and 0.75mM SA equally increased the P_n_ whereas at 50% FC, 1mM SA was effective and at 75% FC, 0.50mM SA increases P_n_ in comparison to respective control. While, at the flowering stage, 1mM, 0.50mM and 0.25mM SA concentration was found equally efficient in increasing P_n_ at 25% FC. Reduction in these photosynthetic parameters was found positively correlated with yield reduction. Similar results were reported by **Liang et al. (2020)** in *Solanum lycopersicum*. There was observed a similar trend for the stomatal conductance at 25% FC where 1mM SA was effective. SA was found equally effective in increasing carboxylation efficiency, water use efficiency and intrinsic water use efficiency. Similarly, an increase in P_n_ and g_s_ were observed in *Fragaria × ananassa* under drought stress by exogenous application of SA **(Ghaderi et al., 2015).** Likewise, seed priming with SA enhances the photosynthesis rate (P_n_), stomatal conductance (g_s_), transpiration rate (T_r_), intercellular CO_2_ (C_i_) and water use efficiency (WUE) in *Carthamus tinctorius* seedlings under drought stress **(Mohammadi et al., 2017).** In *Sesamum indicum*, SA seems to increase P_n_, g_s_ and T_r_ under drought might be due to the role of SA in decreasing ABA levels **(Pourghasemian et al., 2020)**. The water use efficiency of plants was affected negatively by the reduced soil water. In the present study, SA at 1mM concentration for 25% FC, 0.75mM SA concentration for 50% FC as well as for 75% FC and 0.50mM SA concentration at 100% FC increases the WUE during the vegetative stage whereas at the flowering stage, 0.25mM, 0.50mM and 0.75mM SA concentration respectively was found increasing WUE. A similar effect of SA at 1mM as well as 2mM concentration increases the water use efficiency in *Rosmarinus officinalis* under drought stress **(Abbaszadeh et al., 2020)**. Since, SA modulates physiological processes and growth along with the absorption of water. Hence, related findings were reported by **Stallmann et al. (2020)** for the effect of SA on the intrinsic water use efficiency of *Pelargonium x hortorum*.

Cellular membrane stability is an established index to assess crop plants tolerance against abiotic stress. Momentous changes were noted on membrane stability index in both *Cenchrus americanus* and *Triticum aestivum* under drought **(Yadav et al., 2020; Choudhury et al., 2022).** Abiotic stresses especially drought augment the degree of membrane fatty acids saturation by increased permeability of plasma membrane through changing the properties of proteins **(Rawat et al., 2020).** Severe stress condition increases the electrical conductivity that indicates the elevated leakage of ions from leaf corresponding to the increase in membrane injury. Membrane stability index and electrolyte leakage shows negative correlation with the relative water content (Fig.1). In present experiment of *Valeriana*, relative water content (RWC) progressively declines with the increasing water deficit conditions whereas electrolyte leakage and membrane stability index (MSI) seem to increase with the decreasing water levels. The RWC significantly decreases with the increase in permeability of plasma membrane and reduced water supply. The reduction in RWC indicated a loss of turgor that resulted in limited water availability for the cell expansion process and subsequently suppressed plant growth development **(Sharma et al., 2023).** Relative water content (RWC), membrane stability index (MSI) and % conductivity was equally affected by increasing drought. Under water deficit conditions, RWC decreases from 89.95% to 82.26% as the water level drops from 100% FC to 25% FC. RWC can be used to understand the tolerance ability towards water stress in plants. Moreover, MSI decreases from 50.23 to 23.08 with decreases in water levels whereas % conductivity increases with an increase in drought level which might be due to the outflow of mineral ions as a result of cellular membrane damage by reduction in water content in cell and synthesis of ROS. Electrolyte leakage indicates damage to the cell membrane due to impaired membrane integrity **(Yadav et al., 2019).** An increase in the % conductivity was found to be directly correlated with the electrolyte leakage which was the result of protoplasm dehydration causing oxidative stress which was reported by **Raihan et al.(2023)** in *Brassica campestris*and by **Abdelaal et al.(2020)** in *Hordeum vulgare*. RWC under drought decreases by increasing evapotranspiration in plants and reducing the growth of roots **(Sayyari et al., 2013)**.

SA induces plant tolerance towards drought stress by decreasing transpiration rate and stomatal closure. It enhances the rate of photosynthesis by enrichment of intercellular CO_2_ and enhances the nutrient uptake process was observed in spring *Triticum aestivum* under salinity stress **(Alam et al., 2022).** Similarly, SA significantly increases the RWC in plants **(Yousefvand et al., 2022).** SA application maintained the water content of cells up to an optimal level under stress conditions, through the accumulation of osmolytes, which sustained water uptake and increased RWC (**Parveen et al., 2021)**. Similar findings were observed **Estaji & Niknam, (2020)** while working on *Silybum marianum*. *Zea mays* treated with SA show increase in RWC under drought stress than non-treated plants. It might be due to reduced damage to the cellular membrane and a decline in the concentration of ROS **(Shemi et al., 2021).** Similarly, membrane stability decreases with increasing stress as the water level decreases from 100% FC to 25% FC whereas SA increases the membrane stability. Membrane stability index increases in the SA treated plants in comparison to non-treated *Triticum aestivum* **(Khalvandi et al., 2021).** A similar effect of SA was observed by **Ahmad et al. (2017)** in *Pisum sativum* under salinity stress. In *Brassica juncea*, 150 ppm of SA shows a significant increase in MSI by 26.64% during the vegetative stage and 100 ppm during the flowering stage by 13.41% **(Godara et al., 2017)**.Likewise, a study states that a fall off in electrolyte leakage content by means of SA pre-treatment can be related to the amelioration of the antioxidant defence system in the presence of SA under PEG stress **(Torun et al., 2024)**. SA has been observed to mitigate the adverse effect of stresses on membrane integrity through the activation and accumulation of anti-oxidative species which plays a crucial role in sustaining the redox state of membrane proteins **(Ali et al., 2023)**.**Shehzad et al., (2019)** also observed the effectiveness of exogenous SA application in reducing adverse effects of drought stress in *Helianthus annuus*. The increase of growth by SA treatment in drought-stressed plants could be due to the protective role of SA in membrane integrity and regulation of ion uptake. RWC was reduced under drought stress in agreement with results obtained in *Hordeum vulgare* **(Gan et al., 2015),** *Triticum aestivum* **(Barquero et al., 2022)** and *Oryza sativa* **(Dien et al., 2019)**. SA treatment caused increased RWC under drought which was similar to **Antoni**ć **et al. (2020)** observations. SA caused increased accumulation of compatible osmolytes in plants and after that increased water uptake **(Islam et al., 2023).**

In a current study of *Valeriana,* the maximum photochemical efficiency was observed to reduce during water stress conditions whereas, electron transport flux (ET_o_/RC) increases suggesting that drought limits the electron transport to the PS II and causes the closer of the PS II thus reduces the rate of photochemical reaction. F_v_/F_m_ is positively correlated with the photosynthetic rate of plant. This can be used as a rapid and efficient method for the estimation of the drought on plant productivity under drought stress. The similar results for the correlation between the chlorophyll fluorescence and photosynthetic parameters were observed by (**Pappula-Reddy et al., 2024)**. Reduction in the photosynthetic rate and carboxylation efficiency is also evident in the destructive effect of water scarcity on the photosynthetic apparatus. In the current investigation, F_v_/F_m_ declined under the effect of drought stresses which might be due to instability in the membrane integrity and dysfunction of thylakoids in chloroplasts **(Pandey et al., 2022; Manaa et al., 2021)**. During the photosynthesis process in plants, light energy absorbed by chloroplasts is dissipated through three reciprocally related pathways: photosynthetic electron transport, chlorophyll flourescence and heat dissipation. As stress increases the heat dissipation capacity reduces and damages PS II. F_v_/F_m_ reflects the light use efficiency of the plant and represents the efficiency of antenna pigment to transport the absorbed light to the PS II which is then utilized for the photochemical reaction under ambient conditions **(Shanker et al., 2022).** Light absorption and the efficiency of its conversion into chemical energy is considered the most influential eco-physiological tool to study the photosynthetic process in plants **(Li et al., 2023)**. It is the reflective maximum photosynthesis capacity after the optimal dark adaptation **(Ito et al., 2021)**. The fluorescence method is being successfully used in the identification of drought-tolerant lines in *Triticum aestivum* (**Peršić et al., 2022**). In the present study, a drought-stressed plant shows a reduction in the maximum photochemical efficiency of PS II and reduces the quantum yield **(Manaa et al., 2021).** Minimum photochemical efficiency was absorbed as the drought level increased; a significant and gradual decrease was found at 25% FC. Whereas, the increase in electron transport flux per reaction centre of PS II increases with an increase in drought level. A similar reduction in electron flux was recorded in *Solanum lycopersicum* under drought **(Zhang et al., 2017)**. The reduction in the efficiency of PS II directly affects plant growth and yield. Related findings were observed by **Sánchez-Reinoso et al. (2018)** while working on *Phaseolus vulgaris* it might be due to the destruction of chlorophyll pigment and photosynthetic apparatus. A decline in photochemical efficiency reduces the photosynthetic rate and ultimately affects plant growth. Likewise, in drought sensitive genotypes the maximum photochemical efficiency was found lower than drought tolerant genotypes under drought stress **(Badr & Brüggemann, 2020).** In *Triticum aestivum*, the yield was found positively correlated with the chlorophyll fluorescence thus it is a reliable parameter for the evaluation of the SA effect on plants under stress **(Ali et al., 2023).** SA increases the photochemical efficiency by reducing damage to the photosynthetic apparatus with the increasing drought. It was observed that 1mM SA effectively increase photochemical efficiency and balances the electron transport to the PS II in *Vigna radiata* **(Lotfi *et al.,* 2020)**. A similar effect of 1mM SA was reported by **Jahan *et al*. (2019)** in *Solanum lycopersicum* under heat stress. SA in *Vigna radiata* regulates the K^+^ accumulation and improves the activity of PS II under salinity stress than non-SA treated plants. It was also found to improve the balance of electron transport to the PS II **(Ghassemi-Golezani and Lotfi, 2015)**. Increased gas exchange attributed to the application of SA might improve photosynthetic efficiency and chlorophyll fluorescence enabling plants to withstand environmental stresses **(Moustakas et al., 2022)**.

An increase in water scarcity decreases the pigment content (chlorophyll and carotenoid content) in *Valeriana* because of a reduction in water flow that results in less absorption of mineral nutrients such as nitrogen and magnesium which are essential elements in the chlorophyll formation. It results in non-functional thylakoid, photoassimilate level, dehydration of protoplasm and oxidative injury to chloroplasts. It also induces stomatal closure and decreases CO_2_ concentration in the mesophyll cells. Very severe drought conditions cause limited photosynthesis due to a decline in RubisCO activity and reduced gas exchange **(Dubberstein et al., 2020; Hasanagić et al., 2020)**. A decline in chlorophyll pigment under water scarcity either in *Cenchrus americanus* or in *Triticum aestivum* could be ascribed to the rate of chlorophyll degradation or reduction in the activity of chlorophyll biosynthesis enzymes **(Choudhury et al., 2022; Popova et al., 2023).** It was also observed that under drought stress the pigment content in *Mentha arvensis* decreases significantly **(Elhakem, 2019).** Similarly, in *Oryza sativa* drought stress caused a significant decrease in pigment contents: chlorophyll “a”, “b”, a/b, total chlorophyll and carotenoids content **(Nasrin et al., 2020)**. Photosynthesis is a vital plant metabolic pathway. The management of plant growth under drought stress requires upholding the optimum photosynthetic rate **(Zahra et al., 2023)**. Levels of accessory photosynthetic pigments such as carotenoids altered during physiological and pathological conditions. Apart from their light harvesting function and contribution to photosynthesis, carotenoids have a further function in thylakoid lamellae to protect chlorophylls against oxidative annihilation by O_2_ when the irradiance level is high **(Caferri et al., 2022). Zhang et al. (2022)** indicate that higher levels of pigment content (carotenoid and chlorophyll) might prove an effective index for the selection of superior-quality genotypes under saline conditions.

Foliar application of SA minimizes drought stress in *Valeriana*. In the present experiment, it was observed that at 25% FC, 0.75mM SA concentration during the vegetative stage and 1mM SA concentration during the flowering stage effectively promotes the chlorophyll “a”, total chlorophyll and carotenoid content. While 0.50 mM SA was more effective at both stages in chlorophyll “b” for reducing damage. At moderate and mild stress levels (50% FC and 75% FC), 0.25mM and 0.50mM SA concentrations were effective for the enhancement of pigment contents at both vegetative and flowering stages. Similar to our results, the application of SA enhances chlorophyll content in *Aloysia citrodora* indicating the involvement of SA in photosynthesis. Thus, the results obtained explain the positive impact of SA that has increased resistance towards stress conditions by induction of photosynthetic rate **(Khalvandi et al., 2021)**. Also in other studies, the application of SA increases pigment contents in *Glycine max* (**Kaya et al., 2024)**, *Zea mays* **(Shemi et al., 2021)** and *Triticum aestivum* **(Parveen et al., 2021)** cultivated under drought stress conditions. Similarly, **Hafeez et al. (2024)** reported that photosynthetic pigment contents had been reduced significantly under drought stress whereas SA could alleviate this effect by moving the intracellular nitrate resources or increase of pigment biosynthesis. Accordingly, **Parveen et al. (2021)** SA protected photosynthesis and enhanced RubisCO activity in treated wheat under water stress. Chlorophyll is an important pigment molecule associated with photosynthesis hence plant bioregulators like SA maintain cellular osmoticum which aids in enhancing the chlorophyll “a” and “b” contents **(Mandal & Dutta, 2020).** SA decreases the destruction of photosynthetic apparatus and enhances the stability of PS I and PS II. It also stimulates the synthesis of photosynthetic pigment which might be due to the increase in the nutrient uptake production of antioxidant enzymes **(Kusvuran & Yilmaz, 2023; Lobato et al., 2021).**

## 5. Conclusion

Increasing drought stress results in a damaging effect on physiological traits showing a negative effect on yield (Fig. 4). Since, to improve physiological traits foliar spray of SA shows a positive effect in *V. wallichii.* Different doses of SA under various levels of drought stress might be used to overcome the hazardous effects of drought. It was found that at 100% and 75% FC, 0.25mM and 0.50mM SA concentrations were found effective during vegetative as well as flowering stages for most of the traits under study. At 75% FC, 1mM is useful during the vegetative stage. At 50% FC, different concentration of SA positively affects the vegetative stage (*i.e.,* 0.25mM SA concentration), and the flowering stage in leaves (*i.e.,* 0.50mM SA concentration). Similarly, at 25% FC, 1mM concentration is found best among all the concentrations during both the stages, *i.e.* vegetative and flowering stages in comparison to respective control values.

**Fig. 3.**
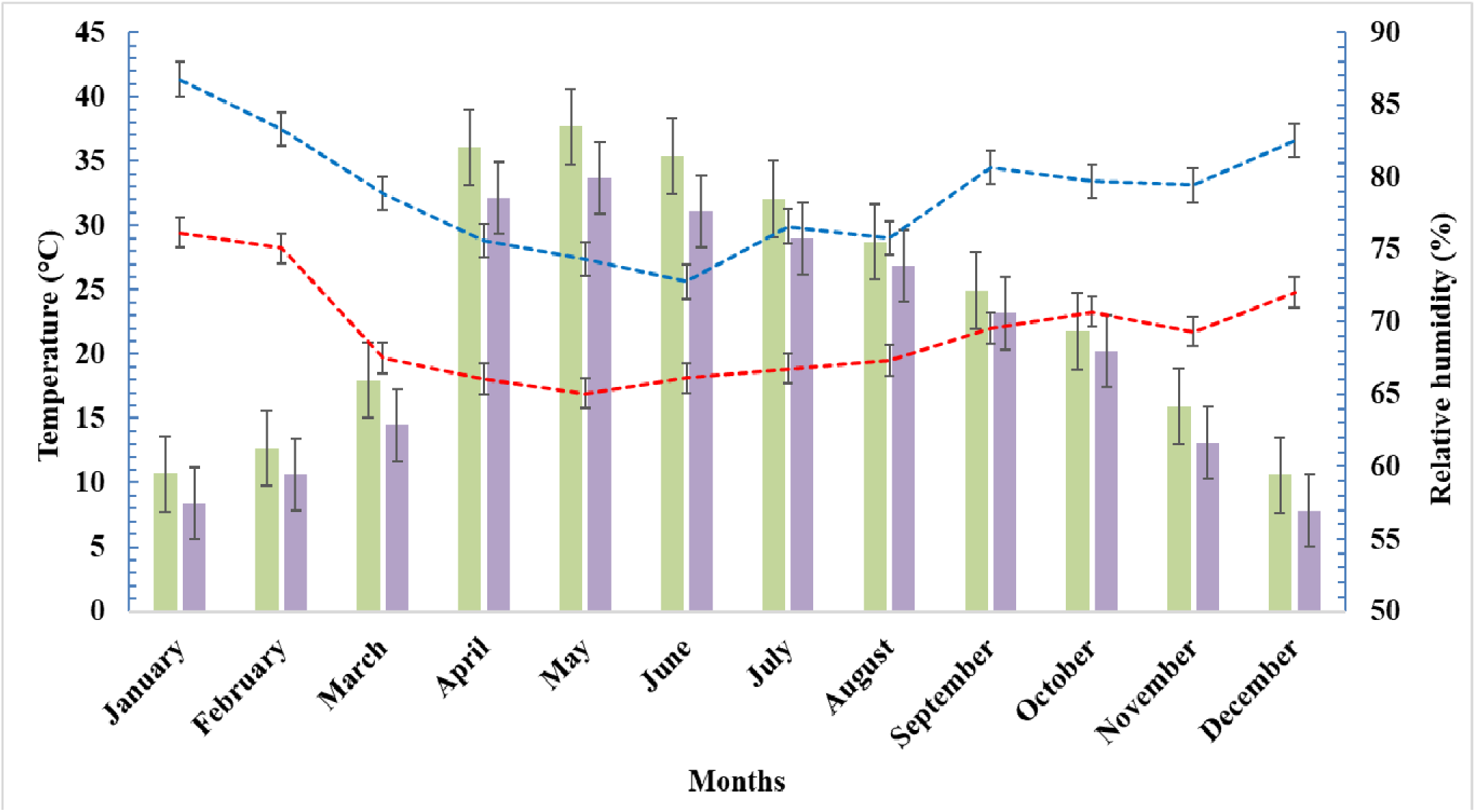
The cumulative weather data, including temperature and relative humidity, recorded under control conditions for the crop seasons 2018-2019 and 2019-2020. The combined data provide a comprehensive overview of the environmental parameters influencing crop performance across both years

**Fig. 4.**
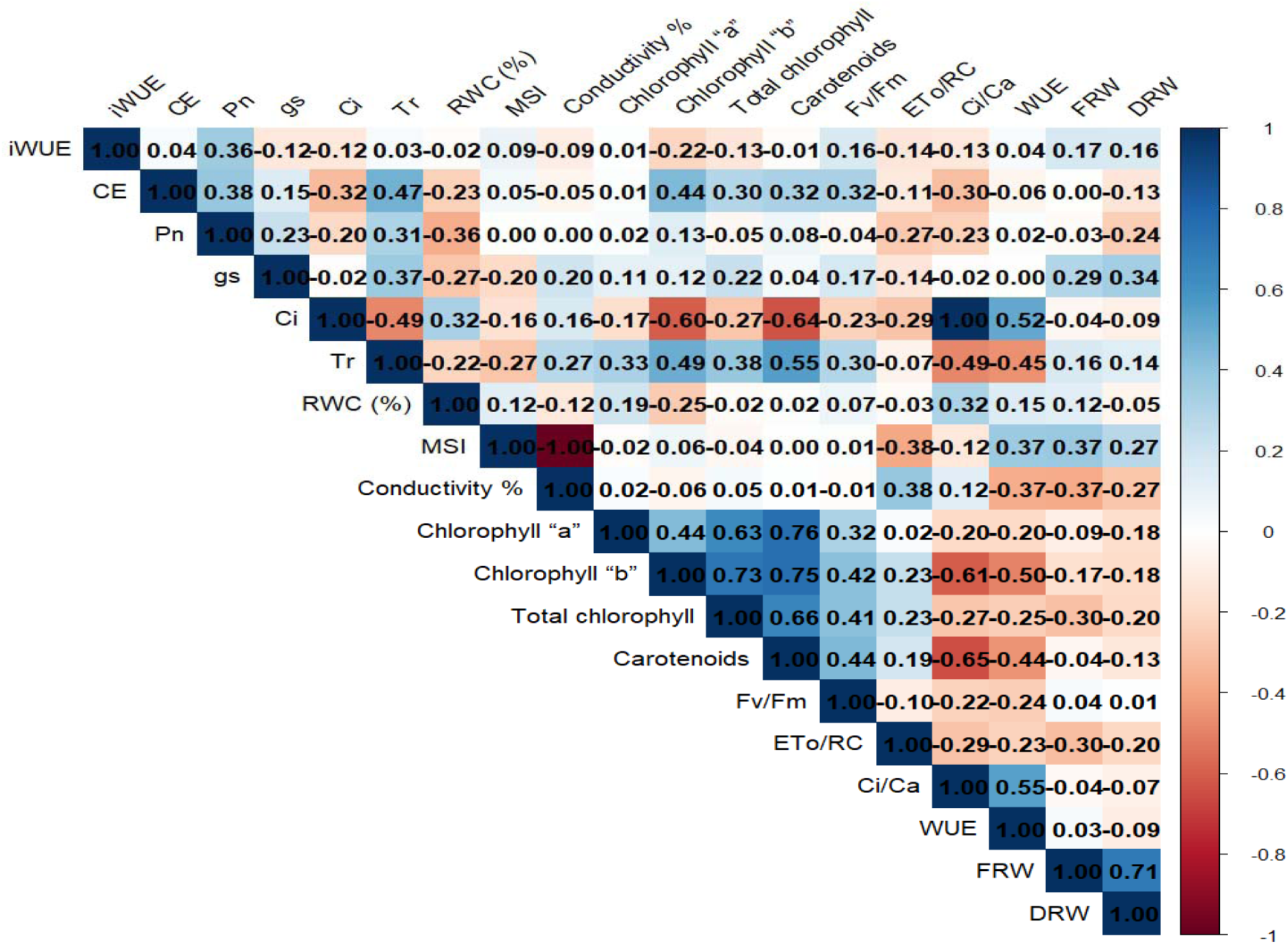
Correlation between different parameters (Darker colours show strong relationships; Lighter colours show weak relationships and numbers show the strength (closer to 1 or -1 means strong correlation). **Note: Fv/Fm-** Maximum photochemical efficiency, **(ETo/RC)-** Electron transport flux/ reaction centre of PS II, **CO_2_ (Ci/Ca)-** Ratio of internal to ambient, **WUE-** Water use efficiency, **iWUE-** Intrinsic water use efficiency, **CE -** Carboxylation efficiency, **Pn-**Photosynthesis rate, **gs-** Stomatal conductance rate, **Ci-** Intercellular CO_2_ concentration, **Tr-**Transpiration rate, **RWC-** Relative water content, **MSI-** Membrane stability index,% **Conductivity-** Percent Conductivity,**FRW-** Fresh root weight, **DRW-** Dry root weight

## Acknowledgement

We would like to thank the Director of High-Altitude Plant Physiology (HAPPRC), HNB Garhwal University, Srinagar Garhwal, Uttarakhand; India.

## Author Contribution Statement

Conceptualization, draft writing, investigation, and methodology done by SA and DJ. Statistical analysis was done by DJ. Review, editing and supervision were done by BP, HCJ & MKB.

